# L1EM: A tool for accurate locus specific LINE-1 RNA quantification

**DOI:** 10.1101/714014

**Authors:** Wilson McKerrow, David Fenyö

**Affiliations:** Institute for Systems Genetics, NYU School of Medicine, 550 1^st^ Ave, New York, NY, USA

## Abstract

**Motivation:** LINE-1 elements are retrotransposons that are capable of copying their sequence to new genomic loci. LINE-1 derepression is associated with a number of disease states, and has the potential to cause significant cellular damage. Because LINE-1 elements are repetitive, it is difficult to quantify RNA at specific LINE-1 loci and to separate transcripts with protein coding capability from other sources of LINE-1 RNA.

**Results:** We provide a tool, L1-EM that uses the expectation maximization algorithm to quantify LINE-1 RNA at each genomic locus, separating transcripts that are capable of generating retrotransposition from those that are not. We show the accuracy of L1-EM on simulated data and against long read sequencing from HEK cells.

**Availability:** L1-EM is written in python. The source code along with the necessary annotations are available at https://github.com/FenyoLab/L1EM and distributed under GPLv3.

## Introduction

Retrotransposons are genomic sequences that are able to copy themselves to a new location via an RNA intermediate. Long Interspersed Element 1 (LINE-1) is the only retrotransposon known to be capable of autonomous retrotransposition in the human genome. LINE-1 retrotransposition begins with transcription from one or more genomic loci. The unspliced LINE-1 RNA is polyadenylated and exported from the nucleus, where two proteins – ORF1p and ORF2p – are translated from separate open reading frames on the LINE-1 RNA. ORF1p is an RNA binding protein (Martin 2006) while ORF2p possesses the endonuclease and reverse transcriptase activity necessary for retrotransposition (Hattori et al. 1986; Feng et al. 1996). The LINE-1 RNA then forms a ribonucleoprotein complex (RNP) with its protein products (Wei et al. 2001). The RNP is imported into the nucleus, likely during the S phase of cell cycle (Mita et al. 2018), where it is reverse transcribed and inserted at a new genomic locus. At least 17%, and probably more, of the human genome is derived from LINE-1 activity. However, in the human genome, only about 100 LINE-1 copies remain capable of retrotransposition. Most of the about 500,000 LINE-1 copies in human genome are either ancient, truncated or have premature stop codons that prevent the translation of intact ORF1p or ORF2p.

In modern humans, LINE-1 activity has been observed in early embryonic development (Kano et al. 2009), during neuronal differentiation (Coufal et al. 2009), in cancer (Rodić et al. 2014; Tubio et al. 2014), and during aging (De Cecco et al. 2019). LINE-1 is theorized to contribute to genomic instability in cancer (Kemp and Longworth 2015), play a regulatory role in stem cells and in early development (Percharde et al. 2018; Rodriguez-Terrones and Torres-Padilla 2018), trigger inflammation in senescent cells (De Cecco et al. 2019) and provide genomic plasticity to neurons (Singer et al. 2010). However, despite a growing appreciation for the ubiquity of LINE-1 RNA, the repetitive nature of LINE-1 sequences makes accurate measurements of LINE-1 RNA difficult to obtain. Many studies look only at a global measure of LINE-1 RNA, such as that provided by a qPCR assay. However, measurements that do not quantify specific LINE-1 transcripts fail to differentiate retrotransposition competent LINE-1 transcripts from LINE-1 transcripts that lack intact open reading frames and from other transcripts, such as long non-coding RNAs, that may passively include some LINE-1 sequence. Recent evidence both at the level of RNA (Philippe et al. 2016; Deininger et al. 2017) and at reinsertion (Tubio et al. 2014) indicate that a small number of loci are responsible for most of the LINE-1 activity, further highlighting the need for locus specific LINE-1 RNA quantification.

In this paper we describe a method, L1-EM, that uses the expectation maximization algorithm (EM) (Dempster et al. 1977) to quantify LINE-1 transcripts from paired-end RNAseq reads. We focus on the newest elements in the human genome: L1HS. This family includes the retrotransposition competent elements and is the most difficult to align reads to. For each L1HS locus that includes the 5’UTR sequence, we quantify proper LINE-1 transcription that begins at the 5’UTR and separate it from passive co-transcription that begins from another transcription start site but includes some LINE-1 sequence. Transcription from older L1PA elements is also quantified, but not discussed here as these elements do not contribute to active retrotransposition.

Several methods have been proposed to quantify LINE-1 locus expression, but none are as complete as the L1-EM method we propose. Philippe et al. (Philippe et al. 2016) measured transcripts that overrun the 3’ end of LINE-1, but our results indicate that the volume of this “run-on” transcription varies between loci (see results, figure 4.) Deininger et al. (Deininger et al. 2017) identified transcribed loci by looking at the subset of LINE-1 reads that align uniquely, but this accounts for only a small fraction of the reads aligning to the young L1HS elements. Other algorithms do exist that use EM to quantify transposable element RNA sequences (eg Jin et al. 2015; Yang et al. 2019), but none account for passive co-transcription.

## Methods

### The transcripts

All LINE-1 loci in the hg38 human reference genome that are labeled as L1HS or L1PA* (i.e. L1PA2, L1PA3, etc.) in RepeatMasker (Bao, Kojima, and Kohany 2015) are divided into two categories. Category 1 includes all elements that include the LINE-1 5’UTR, and Category 0 consists of elements that do not include the 5’UTR. Because the 5’UTR functions as the LINE-1 promoter, only elements in category 1 are allowed to form LINE-1 transcripts. For each element with a 5’UTR (category 1), five transcripts, illustrated in figure 1A, are considered:

1. The “only” transcript that runs sense and includes only the annotated element.
2. The “Run-on” transcript that runs sense and includes downstream sequence.
3. The “Passive” sense transcript that runs sense and includes both upstream and downstream sequence.
4. The “Passive” antisense transcript that runs antisense and includes both upstream and downstream sequence.
5. The “Antisense” transcript that runs antisense and includes only the first 500 bases of the element plus upstream (downstream on the antisense strand) sequence. (Speek 2001)

**Figure 1.**
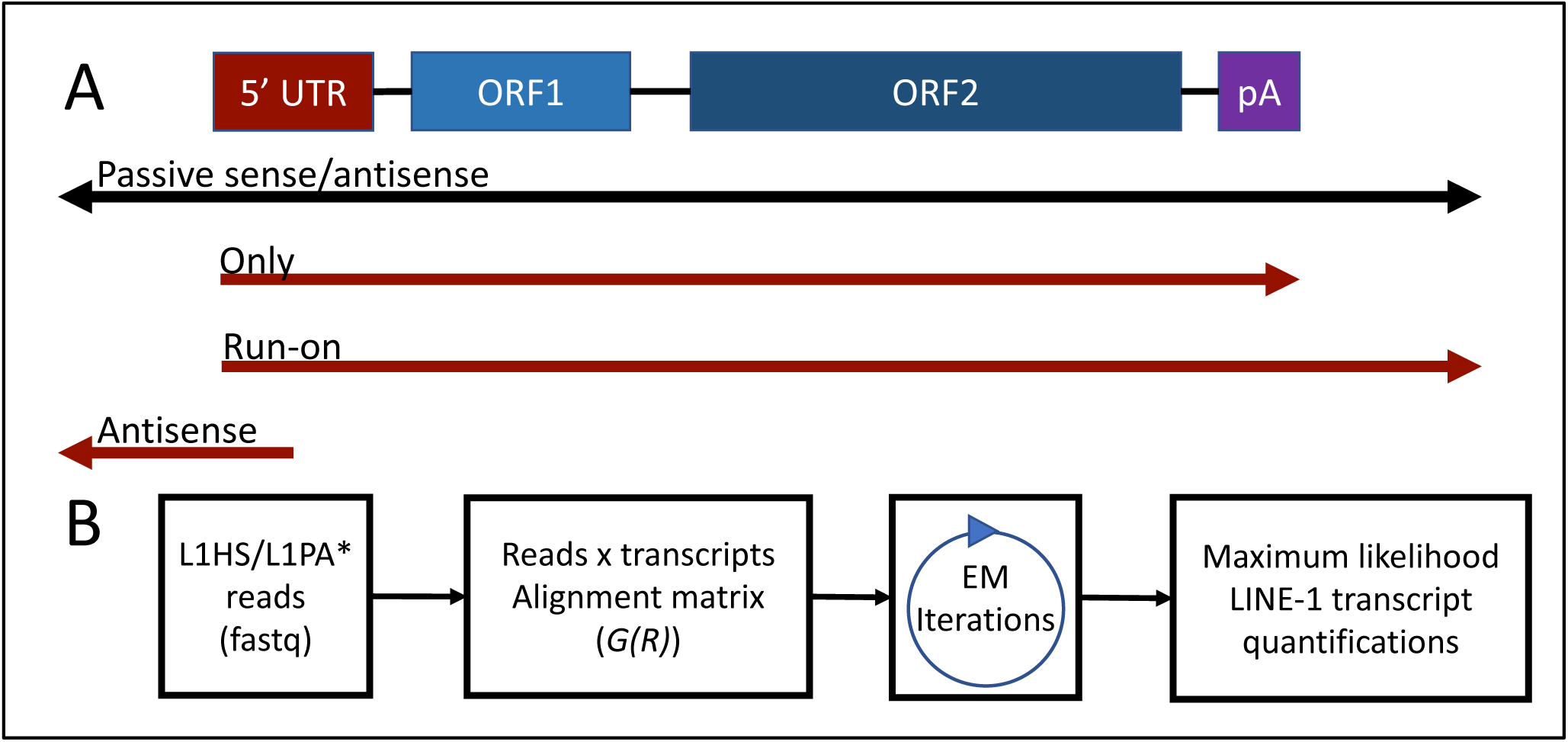
Transcripts and pipeline. (A) Types of transcripts that include LINE-1 sequence. “Run-on”, “only” and “antisense” are only allowed at loci with 5’UTRs. “Passive” transcripts are allowed at all reference loci. (B) Outline of the L1-EM pipeline.

For elements without a 5’UTR (category 0), only the “Passive” transcripts are included. Exon sequences in or within 400 basepairs of an L1HS or L1PA* element are also included to prevent host mRNAs from being conflated with LINE-1 expression.

These LINE-1 transcripts can then be quantified using an EM algorithm method that is similar to the methods used to quantify gene isoforms (e.g. (B. Li and Dewey 2011; Patro et al. 2017)). Care however must be taken as there are many highly similar LINE-1 transcripts, and these transcripts differ at mismatches and indels rather than at splice junctions. The details of our application of the EM algorithm to LINE-1 transcript quantification follow.

### The generative model

List of random variables:

- *X* = *X*_1_ … *X*_*i*_ … *X*_*n*_, where ∑ *X*_*i*_=1, is the relative abundance of each transcript enumerated above.
- *R* = *R*_1_ … *R*_*j*_ … *R*_*m*_ are the read sequences.
- *A* = *A*_1_ … *A*_*j*_ … *A*_*m*_ are the read alignments.

The reads are assumed to be independently generated by first sampling a transcript according to *X*, then choosing a random location in that transcript, and finally introducing mismatch/indel read errors with probability *ϵ* (0.01 by default). This model yields the following likelihood function:

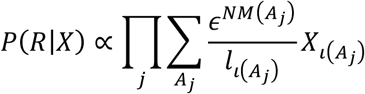

Where *NM* is the edit distance of an alignment, *l*_*i*_ is the effective length of transcript *i*, and *ι*(*A*_*j*_) is the locus that *A*_*j*_ is an alignment to. For “passive” transcripts, the effective length is the length of the element plus the median template length for read pairs. For “only” transcripts, it is the element length minus the median template length. For “run-on” transcripts it is the element length, and for “antisense” transcripts it is 500. Transcripts are given a minimum effective length of 500.

The likelihood function can be simplified by rearranging the sum over *A*_*j*_ to group alignments by transcript:

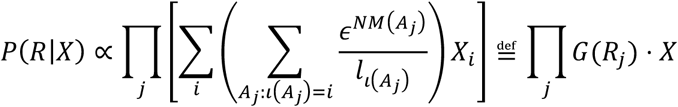

This simplification shows that the likelihood function is a product of linear functions, and is thus convex(Jiang and Wong 2009). We can therefore find the maximum likelihood estimate of the relative expression levels (*X*) using expectation maximization, without risking the identification of local maxima.

### Collection of L1HS/L1PA reads

If reads are not previously aligned to hg38, they are first aligned to hg38 using bwa aln (H. Li and Durbin 2009), allowing an edit distance of up to 3. If reads are aligned, unaligned reads are extracted and realigned to hg38 using bwa aln. This realignment step is necessary as many aligners do not report alignments for highly ambiguous reads. Once genome alignments are complete, any read pair that overlaps an L1HS or L1PA* family element in either the initial alignment or the realignment is extracted for further analysis.

### Generation of candidate alignments

Because it is not feasible to sum over all possible alignments of every read, we need to approximate *G*(*R*_*j*_) by only summing over a set of candidate alignments. This set of alignments is generated by using bwa aln to find all alignments to the transcripts enumerated above that have an edit distance less than or equal to 5, and no indel within 20 basepairs of the end of the read. Candidate alignments that have at most 2 more mismatches than the best alignment for that read are retained. The alignment likelihood, 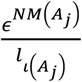, is then calculated for each candidate alignment and added to a *m* × *n* sparse matrix, *G*(*R*), where

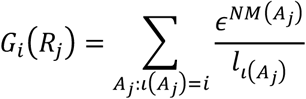

is proportional to the likelihood that a random read from transcript *i* will have the sequence *R*_*j*_.

### Estimation of transcript abundance

The EM algorithm requires iterating between two steps: first, during the E step, an estimate of the relative expression levels, *X*^(*t*)^, is fixed, and the expected number of reads originating from each transcript is calculated. Second, during the M step, a new estimate of the expression level, *X*^(*t*+1)^, is calculated by normalizing the expected counts. This yields the following iteration:

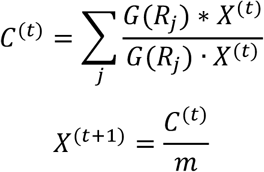

where the * operation indicates elementwise multiplication. Because the likelihood function is a product of linear functions, we can start with any initial guess, *X*^(1)^, that has no 0 entry and iterate these steps until we converge to the maximum likelihood estimate:

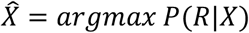

L1-EM uses the uniform distribution as an initial guess, and repeats the iteration until no entry of *X* changes by more than a parameter *δ* (10^−7^ by default). Because some fraction of “passive” transcription can be mislabeled as “only” or “run-on”, “only” and “run-on” transcripts are not reported if they make up less than ¾ of the expression at that locus.

### Implementation

L1-EM is implemented in python, with the pysam library used to read bam format alignment files and the scipy sparse matrix library used for all intensive computations. L1-EM is available as an open source program under the GPLv3 license. Currently only annotations to quantify LINE-1 elements in the hg38 human genome are available, but instructions for generating new annotations for other elements in other genomes are provided.

## Results

### On simulated data, L1-EM provides accurate estimates where the accuracy depends on read coverage

For each L1HS family element with a 5’UTR, an independent value was sampled from an exponential distribution with mean 1. These values were then cubed and normalized to generate a relative expression vector, X. The simulation strategy yields a few highly expressed elements, similar to what has been previously reported (Philippe et al. 2016; Deininger et al. 2017). 30,000 100 base-pair paired end reads were then simulated from the “only” transcript according to the generative model described above. Errors were introduced at each base with probability 0.01. In addition, 250,000 reads were sampled completely at random from L1HS, L1PA* and 400 bases of flanking sequence to simulate noise. Simulated reads were analyzed using the L1-EM pipeline, and the resulting estimates were compared to the simulated read counts (excluding the 250,000 “noise” reads.)

When L1-EM is given all 30,000 L1HS “only” transcript reads, it provides an extremely accurate result. Only about 5% of the total read mass would need to be moved between elements to yield an answer that is exactly correct (figure 2, top right.) As more of the reads are withheld from the pipeline, accuracy weakens: when 10,000 reads are considered, about 10% of the read mass is placed incorrectly; with 5,000 reads, about 15% of the read mass is misplaced; when only 2,000 of the 30,000 reads are considered, about 30% of the read mass would need to be moved to get the simulated expression values. However, when coverage is low, most of the estimation error comes from the elements expressed at a low level. Even with only 2,000 reads, the highly expressed elements are correctly identified and accurately quantified (figure 2, bottom right.)

**Figure 2.**
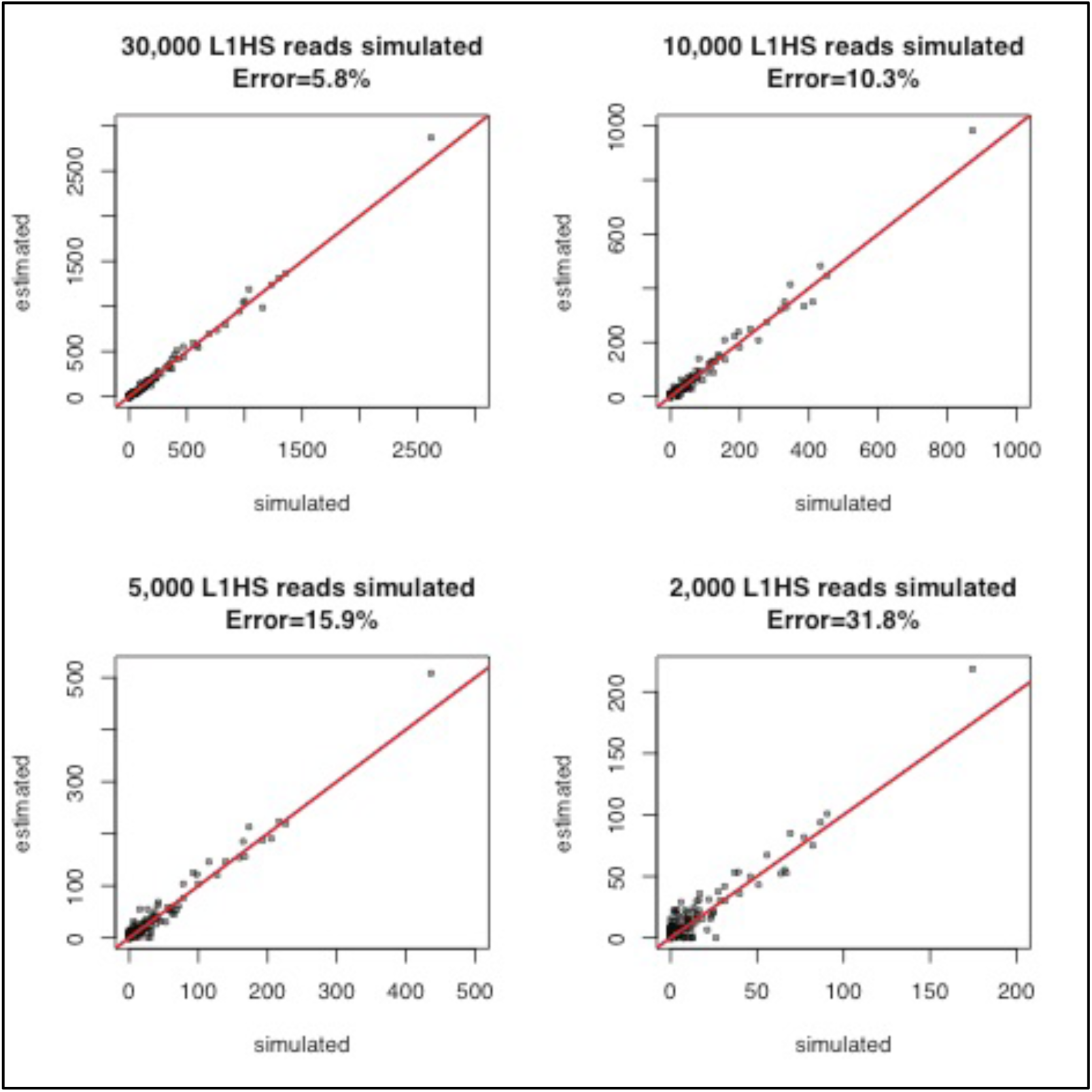
Simulated data. L1EM performs well on simulated data, especially with higher read coverage. The displayed error values are the fraction of reads that would need to be moved to different elements to exactly recoup the simulated expression values.

### L1-EM estimates agree with L1 5’RACE long read sequencing in HEK293 cells

Deininger et al. (Deininger et al. 2017) used the 5’RACE technique to specifically amplify LINE-1 transcripts beginning with the L1 5’UTR in HEK293 cells. PacBio reads exceeding 1000 letters in length were sequenced from the first 1237 bases of the amplified transcripts. We accessed these reads from the SRA database (SRR4099955), aligned them to hg38 using bwa mem, and then counted unique alignments to quantify LINE-1 transcripts beginning with the L1 5’UTR. Deininger et al. did not perform Illumina RNA-seq from these cells, so we ran L1-EM on 150 bp strand-specific HEK293 RNA-seq reads from another study (Aktaş et al. 2017) (SRR3997504-7). To match the L1 5’RACE data, we pooled “only” and “run-on” transcripts.

Only about 2000 reads were estimated to come from L1HS “only” or “run-on” transcripts in this dataset. Nevertheless, L1-EM is able to accurately quantify the two highly expressed elements in HEK293 (figure 3, top left). This quantification remains accurate even when we ignore the strand specifity or trim the read length to 50bp (figure 3, other panels.) However, without strand specificity some elements with no evidence of expression in L1 5’RACE appear to be marginally expressed.

**Figure 3.**
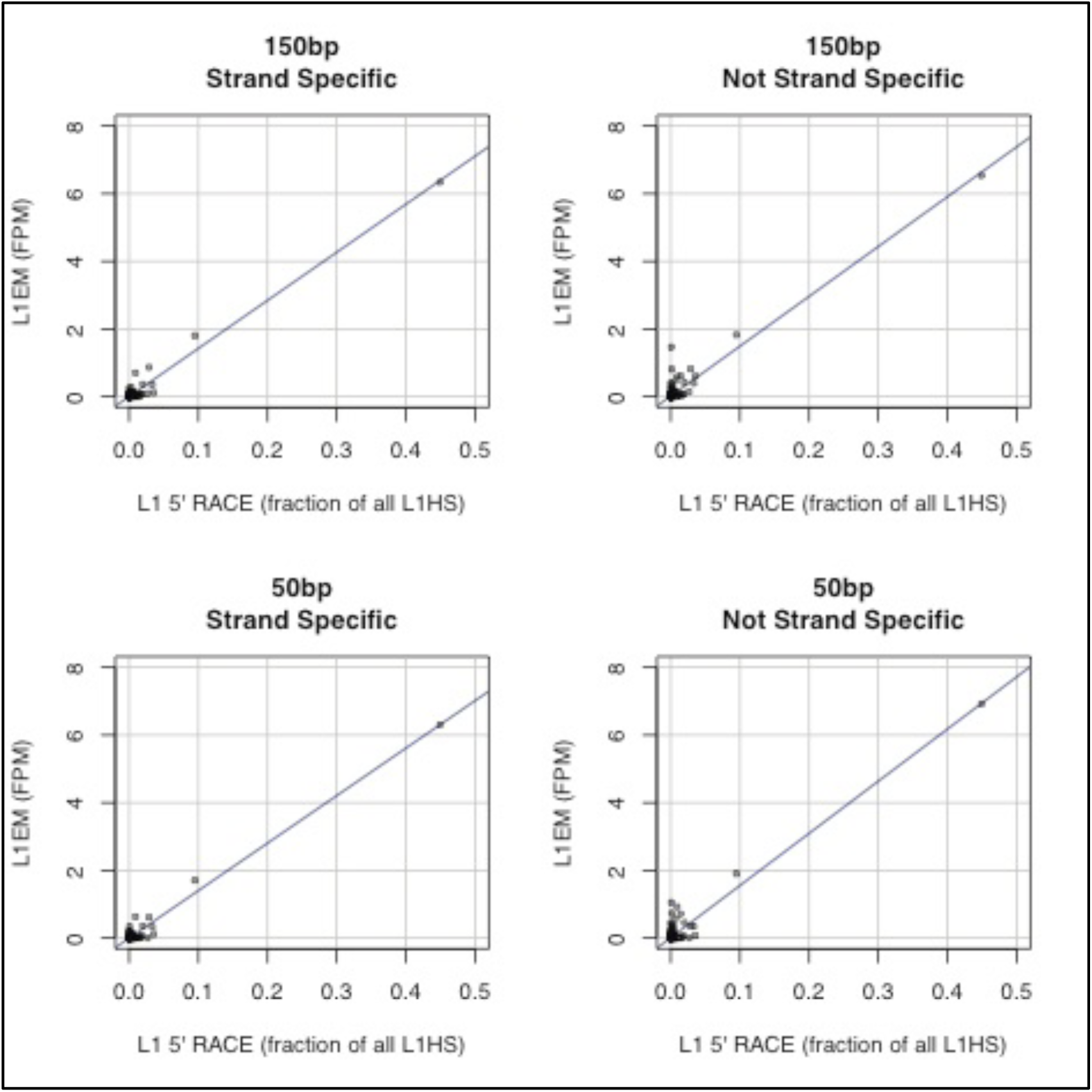
Biological data. L1-EM shows good agreement with long read L1 5’RACE sequencing in HEK293. It is still possible to identify the top expressed elements even if the reads are shortened to 50bp and strand specificity is ignored.

### 3.1 The fraction of transcripts that “run-on” varies considerably and depends on polyA length

A previous study of locus specific LINE-1 RNA, focused only on reads that run through the 3’ polyA tail (“run-on” in our nomenclature) (Philippe et al. 2016). To determine whether LINE-1 transcription can be accurately gauged by considering only run-on reads, we reanalyzed 150bp paired end strand specific RNA-seq data from this study (ERR973734) using L1-EM. L1-EM estimates that there are about 50,000 reads in this dataset estimated to originate from proper (“only” or “run-on”) LINE-1 transcripts.

At many loci we find “only” transcription without “run-on” transcription (figure 4). This is likely due to variation in the 3’ polyA sequence that strengthens or weakens the polymerase II termination signal. Considering the top 50 expressed LINE-1 loci and looking for a window at the 3’ end of each element that is at least 90% A, we find that loci with “run-on” sequence have an average polyA window of length 22.8, while loci that do not “run-on” have an average polyA window of length 36.4 (t-test p<0.0003). Thus, an accurate quantification of LINE-1 loci requires the consideration of LINE-1 transcripts that do not overrun their 3’ end.

**Figure 4.**
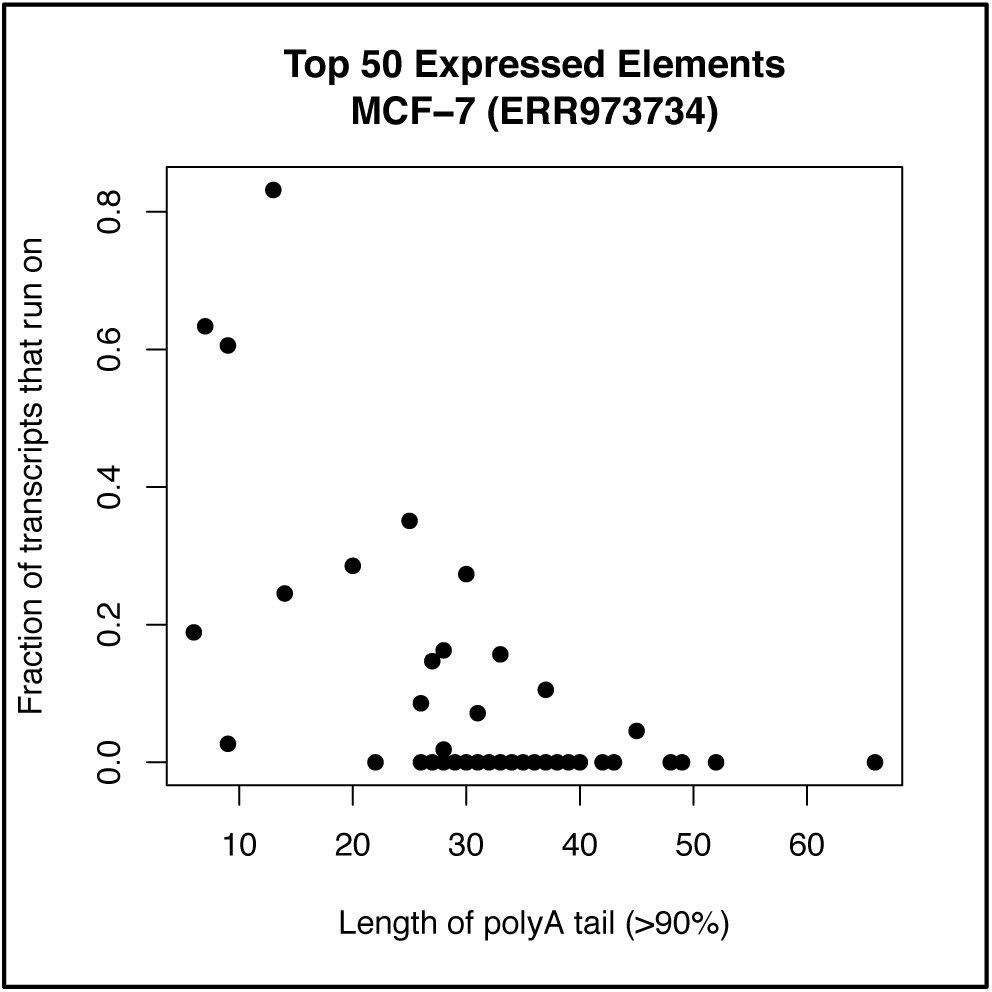
Run-on fraction depends on polyA strength. Fraction of transcripts that run on past the polyA termination sequence varies by element and depends on the length of the polyA sequence.

### 3.2 While the LINE-1 loci expressed vary between samples, one locus at chr22q12.1 tends to be highly expressed

We applied L1EM to all human 100bp paired end, strand specific polyA RNAseq experiments in the ENCODE database (127 datasets including 49 from cell-lines, 16 from in vitro differentiated cells and 62 from tissues.) The full results of this analysis can be found in supplementary table S1. Across these samples, H1 human embryonic stem cells (37 read pairs per million), the MCF-7 breast cancer cell-line (11 FPM), and H1 hESC differentiated into mesendoderm cells (41 FPM) show the greatest expression of L1HS elements with intact ORF1 and ORF2 open reading frames, as annotated in l1base2 (Penzkofer et al. 2017). The chr22:28663283-28669315 locus (chr22q12.1), located in the tetratricopeptide repeat gene TTC28, is the most highly expressed locus in H1 (14% of all intact L1HS expression) and MCF-7 (58%) and is the second most highly expressed in mesendoderm (8%). Across all 127 samples, this locus accounts for 17% of all intact L1HS RNA expression, more than double any other locus.

We find L1HS expression to be low in healthy human tissues, reflecting the widespread assumption that LINE-1 is silenced in somatic tissue. However, we do find expression at at least 1 read pair per million in the brain (1.4 FPM), the esophagus (3.5 FPM), the heart (1.2 FPM), the ovaries (1.4 FPM), the testis (1.7 FPM) and the thymus (1.8 FPM). In the brain (70%), the esophagus (28%) and the thymus (16%), the chr22q12.1 locus is the most highly expressed intact L1HS element. In the heart, the chr4:21159390-21165421 locus located in the voltage-gated potassium channel interacting protein KCNIP4 accounts for 19% of intact L1HS expression. In ovaries and testis, we do not find good agreement between the loci expressed in different samples.

### Measuring total L1HS RNA reveals red herrings

To show the importance of locus specific quantification, we also quantified total L1HS aligning RNA (i.e. including passive transcripts) in the 127 datasets described above. From the total RNA quantifications, some samples appear to express LINE-1, but our previous results indicate that they have little intact L1HS transcription. For example, both samples of H1 differentiated into neurons (ENCFF679IPE and ENCFF309BMG) show evidence of significant L1HS RNA (at 23 and 29 FPM), but we find that over 80% of this RNA is derived from passive transcription, and less than 5% is derived from intact elements. In MCF-7 and H1 cells, over 90% of the total L1HS RNA is derived from “only” or “run-on” transcripts. A similar situation occurs in left and right ventricle tissue (ENCFF466PKR and ENCFF684RZI), where total L1HS RNA is estimated at 22 and 20 FPM respectively, but only about 20% is derived from proper LINE-1 transcripts.

## Discussion

If expressed, LINE-1 elements are capable of causing significant cellular damage, through retrotransposition (Miki et al. 1992), through endonuclease induced DNA damage (Gasior et al. 2006), or through the induction of autoimmunity (Crow 2010). For this reason, retrotransposons are silenced in healthy somatic tissues, although they are expressed in stem cells (Garcia-Perez et al. 2007) and potentially in the brain (Coufal et al. 2009; Muotri et al. 2010). Because of their potential to cause significant harm, when derepressed, LINE-1 elements are likely to contribute to disease phenotype. However, these sources of cellular damage depend specifically on the expression of intact LINE-1 elements. Because older, inactive copies vastly out number young, intact elements, an increase in LINE-1 RNA may not be due to the expression of intact elements. It may instead result from the transcription of elements with damaged open reading frames or from the passive incorporation of LINE-1 sequence into other transcripts. Indeed, our results show that some samples have significant passive transcription of L1HS but little L1HS specific transcription. Thus, understanding which LINE-1 elements are transcribed and at what level, is a key first step to predicting the role that an increase in LINE-1 RNA is likely to play. We provide a tool, L1-EM that can accurately perform this locus specific quantification.

L1-EM provides accurate estimates of RNA expression from young L1HS LINE-1 loci. It does this by combining maximum likelihood expectation maximization methods with our pre-existing knowledge of LINE-1 transcription: namely (1) that it can only occur from elements that retain the 5’UTR sequence, (2) that it sometimes runs beyond the 3’ polyA sequence and (3) that LINE-1 aligning RNA reads can be generated passively from other transcription not related to LINE-1 activity. Here we show that L1-EM performs accurately on simulated data (figure 2) and that it provides good agreement with L1 5’ RACE pacbio sequencing (figure 3).

Previous methods have been developed to address locus-specific LINE-1 expression, but none are as comprehensive as L1-EM. Philippe et al. (Philippe et al. 2016) measured LINE-1 locus specific expression by identifying an H3K4 methylation signal upstream of an element and a run-on signal downstream of the element. Not only does this method rely on the availability of H3K4me3 CHIP-seq, it also will not provide accurate locus quantifications as the fraction of transcripts that run-on varies widely (figure 4). Deininger et al. (Deininger et al. 2017) identified specifically expressed LINE-1 loci using the subset of reads that align uniquely to a particular LINE-1 element. However, because the intact L1HS LINE-1 elements are highly repetitive, few of the reads aligning to these elements will align uniquely, making accurate quantification impossible. Finally, several methods apply expectation maximization to transposable elements (including LINE-1) but only SQuIRE (Yang et al. 2019) provides locus specific estimates, and none differentiate proper LINE-1 transcripts from passive transcription.

It should be noted that L1-EM makes the implicit assumption that LINE-1 RNA is derived from unspliced copies of reference LINE-1 loci. Individuals can have dozens of polymorphic LINE-1 insertions (Gardner et al. 2017) that may be expressed. RNA reads arising from such expression would be mapped to one of the existing LINE-1 elements, most likely the parent element, as that element will bear the greatest sequence similarity. It is possible that for some of the expressed loci that lack run-on transcription, it is actually a non-reference locus that is expressed. However, because we find a strong relation between the polyA sequence strength and run-on transcription, we believe this is the exception rather than the rule. Because LINE-1 RNAs are not spliced during their lifecycle, we do not expect the lack of splicing to be a significant problem for L1-EM. However, splicing of LINE-1 transcripts has been proposed as a regulatory mechanism (Belancio et al. 2010), so a careful quantification of spliced LINE-1 transcripts is potentially a valuable next step for L1-EM.

## Conclusion

L1-EM uses the expectation maximization algorithm to quantify LINE-1 transcripts arising from the 5’ UTR of each L1HS locus. It provides accurate locus specific quantifications both from simulated data and from real data. L1-EM makes it possible to specifically measure intact LINE-1 transcripts that are (potentially) retrotransposition competent, and to identify critical loci.

## Acknowledgements

We would like to acknowledge Dr. Jef Boeke for providing insight into the mechanisms of LINE-1 transcription and the potential sources of LINE-1 RNA.

## References

Aktaş, Tuğçe, İbrahim Avşar Ilık, Daniel Maticzka, Vivek Bhardwaj, Cecilia Pessoa Rodrigues, Gerhard Mittler, Thomas Manke, Rolf Backofen, and Asifa Akhtar. (2017) DHX9 Suppresses RNA Processing Defects Originating from the Alu Invasion of the Human Genome. Nature 544 (7648): 115–19. https://doi.org/10.1038/nature21715.

Bao, Weidong, Kenji K. Kojima, and Oleksiy Kohany. (2015) Repbase Update, a Database of Repetitive Elements in Eukaryotic Genomes. Mobile DNA 6: 11. https://doi.org/10.1186/s13100-015-0041-9.

Belancio, Victoria P., Astrid M. Roy-Engel, Radhika R. Pochampally, and Prescott Deininger. (2010) Somatic Expression of LINE-1 Elements in Human Tissues. Nucleic Acids Research 38 (12): 3909–22. https://doi.org/10.1093/nar/gkq132.

Coufal, Nicole G., José L. Garcia-Perez, Grace E. Peng, Gene W. Yeo, Yangling Mu, Michael T. Lovci, Maria Morell, K. Sue O’Shea, John V. Moran, and Fred H. Gage. (2009) L1 Retrotransposition in Human Neural Progenitor Cells. Nature 460 (7259): 1127–31. https://doi.org/10.1038/nature08248.

Crow, Mary K. (2010) Long Interspersed Nuclear Elements (LINE-1): Potential Triggers of Systemic Autoimmune Disease. Autoimmunity 43 (1): 7–16. https://doi.org/10.3109/08916930903374865.

De Cecco, Marco, Takahiro Ito, Anna P. Petrashen, Amy E. Elias, Nicholas J. Skvir, Steven W. Criscione, Alberto Caligiana, et al. (2019) L1 Drives IFN in Senescent Cells and Promotes Age-Associated Inflammation. Nature 566 (7742): 73. https://doi.org/10.1038/s41586-018-0784-9.

Deininger, Prescott, Maria E. Morales, Travis B. White, Melody Baddoo, Dale J. Hedges, Geraldine Servant, Sudesh Srivastav, et al. (2017) A Comprehensive Approach to Expression of L1 Loci. Nucleic Acids Research 45 (5): e31. https://doi.org/10.1093/nar/gkw1067.

Dempster, A. P., N. M. Laird, and D. B. Rubin. (1977) Maximum Likelihood from Incomplete Data Via the EM Algorithm. Journal of the Royal Statistical Society: Series B (Methodological) 39 (1): 1–22. https://doi.org/10.1111/j.2517-6161.1977.tb01600.x.

Feng, Q., J. V. Moran, H. H. Kazazian, and J. D. Boeke. (1996) Human L1 Retrotransposon Encodes a Conserved Endonuclease Required for Retrotransposition. Cell 87 (5): 905–16.

Garcia-Perez, Jose L., Maria C. N. Marchetto, Alysson R. Muotri, Nicole G. Coufal, Fred H. Gage, K. Sue O’Shea, and John V. Moran. (2007) LINE-1 Retrotransposition in Human Embryonic Stem Cells. Human Molecular Genetics 16 (13): 1569–77. https://doi.org/10.1093/hmg/ddm105.

Gardner, Eugene J., Vincent K. Lam, Daniel N. Harris, Nelson T. Chuang, Emma C. Scott, William S. Pittard, Ryan E. Mills, 1000 Genomes Project Consortium, and Scott E. Devine. (2017) The Mobile Element Locator Tool (MELT): Population-Scale Mobile Element Discovery and Biology. Genome Research, August, gr.218032.116. https://doi.org/10.1101/gr.218032.116.

Gasior, Stephen L., Timothy P. Wakeman, Bo Xu, and Prescott L. Deininger. (2006) The Human LINE-1 Retrotransposon Creates DNA Double-Strand Breaks. Journal of Molecular Biology 357 (5): 1383–93. https://doi.org/10.1016/j.jmb.2006.01.089.

Hattori, Masahira, Satoru Kuhara, Osamu Takenaka, and Yoshiyuki Sakaki. (1986) L1 Family of Repetitive DNA Sequences in Primates May Be Derived from a Sequence Encoding a Reverse Transcriptase-Related Protein. Nature 321 (6070): 625. https://doi.org/10.1038/321625a0.

Jiang, Hui, and Wing Hung Wong. (2009) Statistical Inferences for Isoform Expression in RNA-Seq. Bioinformatics 25 (8): 1026–32. https://doi.org/10.1093/bioinformatics/btp113.

Jin, Ying, Oliver H. Tam, Eric Paniagua, and Molly Hammell. (2015) TEtranscripts: A Package for Including Transposable Elements in Differential Expression Analysis of RNA-Seq Datasets. Bioinformatics (Oxford, England) 31 (22): 3593–99. https://doi.org/10.1093/bioinformatics/btv422.

Kano, Hiroki, Irene Godoy, Christine Courtney, Melissa R. Vetter, George L. Gerton, Eric M. Ostertag, and Haig H. Kazazian. (2009) L1 Retrotransposition Occurs Mainly in Embryogenesis and Creates Somatic Mosaicism.” Genes & Development 23 (11): 1303–12. https://doi.org/10.1101/gad.1803909.

Kemp, Jacqueline R., and Michelle S. Longworth. (2015) Crossing the LINE Toward Genomic Instability: LINE-1 Retrotransposition in Cancer.” Frontiers in Chemistry 3 (December). https://doi.org/10.3389/fchem.2015.00068.

Li, Bo, and Colin N. Dewey. (2011) RSEM: Accurate Transcript Quantification from RNA-Seq Data with or without a Reference Genome. BMC Bioinformatics 12 (August): 323. https://doi.org/10.1186/1471-2105-12-323.

Li, Heng, and Richard Durbin. (2009) Fast and Accurate Short Read Alignment with Burrows-Wheeler Transform. Bioinformatics (Oxford, England) 25 (14): 1754–60. https://doi.org/10.1093/bioinformatics/btp324.

Martin, Sandra L. (2006) The ORF1 Protein Encoded by LINE-1: Structure and Function During L1 Retrotransposition. Journal of Biomedicine and Biotechnology 2006. https://doi.org/10.1155/JBB/2006/45621.

Miki, Y., I. Nishisho, A. Horii, Y. Miyoshi, J. Utsunomiya, K. W. Kinzler, B. Vogelstein, and Y. Nakamura. (1992) Disruption of the APC Gene by a Retrotransposal Insertion of L1 Sequence in a Colon Cancer. Cancer Research 52 (3): 643–45.

Mita, Paolo, Aleksandra Wudzinska, Xiaoji Sun, Joshua Andrade, Shruti Nayak, David J. Kahler, Sana Badri, et al. (2018) LINE-1 Protein Localization and Functional Dynamics during the Cell Cycle. ELife. January 8, 2018. https://doi.org/10.7554/eLife.30058.

Muotri, Alysson R., Maria C. N. Marchetto, Nicole G. Coufal, Ruth Oefner, Gene Yeo, Kinichi Nakashima, and Fred H. Gage. (2010) L1 Retrotransposition in Neurons Is Modulated by MeCP2. Nature 468 (7322): 443–46. https://doi.org/10.1038/nature09544.

Patro, Rob, Geet Duggal, Michael I. Love, Rafael A. Irizarry, and Carl Kingsford. (2017) Salmon Provides Fast and Bias-Aware Quantification of Transcript Expression. Nature Methods 14 (4): 417–19. https://doi.org/10.1038/nmeth.4197.

Penzkofer, Tobias, Marten Jäger, Marek Figlerowicz, Richard Badge, Stefan Mundlos, Peter N. Robinson, and Tomasz Zemojtel. (2017) L1Base 2: More Retrotransposition-Active LINE-1s, More Mammalian Genomes. Nucleic Acids Research 45 (D1): D68–73. https://doi.org/10.1093/nar/gkw925.

Percharde, Michelle, Chih-Jen Lin, Yafei Yin, Juan Guan, Gabriel A. Peixoto, Aydan Bulut-Karslioglu, Steffen Biechele, Bo Huang, Xiaohua Shen, and Miguel Ramalho-Santos. (2018) A LINE1-Nucleolin Partnership Regulates Early Development and ESC Identity. Cell 174 (2): 391-405.e19. https://doi.org/10.1016/j.cell.2018.05.043.

Philippe, Claude, Dulce B. Vargas-Landin, Aurélien J. Doucet, Dominic van Essen, Jorge Vera-Otarola, Monika Kuciak, Antoine Corbin, Pilvi Nigumann, and Gaël Cristofari. (2016) Activation of Individual L1 Retrotransposon Instances Is Restricted to Cell-Type Dependent Permissive Loci. ELife 5. https://doi.org/10.7554/eLife.13926.

Rodić, Nemanja, Reema Sharma, Rajni Sharma, John Zampella, Lixin Dai, Martin S. Taylor, Ralph H. Hruban, et al. (2014) Long Interspersed Element-1 Protein Expression Is a Hallmark of Many Human Cancers. The American Journal of Pathology 184 (5): 1280–86. https://doi.org/10.1016/j.ajpath.2014.01.007.

Rodriguez-Terrones, Diego, and Maria-Elena Torres-Padilla. (2018) Nimble and Ready to Mingle: Transposon Outbursts of Early Development. Trends in Genetics: TIG 34 (10): 806–20. https://doi.org/10.1016/j.tig.2018.06.006.

Singer, Tatjana, Michael J. McConnell, Maria C. N. Marchetto, Nicole G. Coufal, and Fred H. Gage. (2010) LINE-1 Retrotransposons: Mediators of Somatic Variation in Neuronal Genomes? Trends in Neurosciences 33 (8): 345–54. https://doi.org/10.1016/j.tins.2010.04.001.

Speek, M. (2001) Antisense Promoter of Human L1 Retrotransposon Drives Transcription of Adjacent Cellular Genes. Molecular and Cellular Biology 21 (6): 1973–85. https://doi.org/10.1128/MCB.21.6.1973-1985.2001.

Tubio, Jose M. C., Yilong Li, Young Seok Ju, Inigo Martincorena, Susanna L. Cooke, Marta Tojo, Gunes Gundem, et al. (2014) Mobile DNA in Cancer. Extensive Transduction of Nonrepetitive DNA Mediated by L1 Retrotransposition in Cancer Genomes. Science (New York, N.Y.) 345 (6196): 1251343. https://doi.org/10.1126/science.1251343.

Wei, Wei, Nicolas Gilbert, Siew Loon Ooi, Joseph F. Lawler, Eric M. Ostertag, Haig H. Kazazian, Jef D. Boeke, and John V. Moran. (2001) Human L1 Retrotransposition: Cis Preference versus Trans Complementation. Molecular and Cellular Biology 21 (4): 1429–39. https://doi.org/10.1128/MCB.21.4.1429-1439.2001.

Yang, Wan R., Daniel Ardeljan, Clarissa N. Pacyna, Lindsay M. Payer, and Kathleen H. Burns. (2019) SQuIRE Reveals Locus-Specific Regulation of Interspersed Repeat Expression. Nucleic Acids Research, January. https://doi.org/10.1093/nar/gky1301.

